# Evaluating the Transferability of Pathology Foundation Models Across Cancer-related H&E Neurodegeneration-related Immunohistochemical Classification Tasks

**DOI:** 10.64898/2026.09.23.753850

**Authors:** Juan C. Vizcarra, Bhavesh C. Kataria, Chen-Nee Chuah, Thomas M. Pearce, Brittany N. Dugger, Carlos A. Candano, Aaron M. Rosado, Lee A. Cooper, David A. Gutman

**Affiliations:** Emory University, Department of Pathology, GA, USA; University of California Davis, Department of Electrical and Computer Engineering, CA, USA; University of Pittsburgh, Department of Pathology, PA, USA; University of California Davis Health, Department of Pathology and Laboratory Medicine, CA, USA; Northwestern University, Department of Pathology, IL, USA

## Abstract

Foundation models (FMs) have rapidly become dominant in artificial intelligence and are increasingly being adopted in computational pathology. Numerous pathology-specific FMs have been developed and evaluated for a variety of downstream tasks, most commonly using frozen image embeddings with linear probes. While a pathology FM, at least implicitly suggests broad reusability across tasks, the transferability of these models to neurodegenerative disease-related tasks remains largely unexplored.

In this study, we evaluated fourteen frozen feature extractors, including general-vision and pathology FMs, across four pathology image classification datasets spanning two specialized neurodegenerative disease immuno-histochemistry (IHC) tasks (Tau neurofibrillary tangle (NFT) and amyloid-β plaque classification) and two cancer-related hematoxylin and eosin (H&E) tasks (the breast tissue BACH dataset and the multi-tissue TIL dataset). The cancer-related H&E datasets represent tasks more closely aligned with the predominant pretraining domain of many current pathology FMs, whereas the neurodegenerative IHC datasets represent a more specialized domain with limited representation in existing FM pretraining cohorts. Frozen embedding linear probes were compared against a conventional supervised ResNet-50 convolutional neural network (CNN) trained directly on image tiles using identical train, validation, and test splits.

Across both neurodegenerative IHC datasets, the supervised CNN substantially outperformed all frozen FM linear probes. In contrast, pathology FMs achieved performance comparable to, and in some cases exceeding, the supervised CNN across the two cancer-related H&E datasets, with UNI2-h achieving the highest performance on BACH and several pathology FMs performing on par with the CNN on the larger TIL dataset. Furthermore, a supervised CNN trained using only 1% of the Tau NFT training data (1,938 tiles) still exceeded the performance of the best frozen FM linear probe trained on the complete dataset. Together, these results suggest the transferability of frozen pathology FMs may depend strongly on how well the downstream task is represented by their pretraining domain.

These findings demonstrate frozen pathology FMs transfer effectively to the cancer-related H&E tasks evaluated here but may be less effective for specialized neurodegenerative IHC tasks less well represented in current FM pretraining cohorts. Expanding pathology FM pretraining datasets to include a broader range of disease domains and staining modalities may therefore improve transferability to specialized pathology applications. Our findings also highlight the continued importance of conventional supervised learning and motivate future work investigating nonlinear probes and end-to-end FM fine-tuning.

## 1. Introduction

Digital pathology has transformed tissue analysis by enabling the digitization of histological slides into high-resolution whole-slide images (WSIs). These images capture tissue morphology at cellular and even subcellular resolution, allowing detailed examination of structures such as individual cell nuclei. However, this level of detail comes at the cost of large image sizes, with individual WSIs often reaching several gigabytes [1]. As the adoption of digital pathology has accelerated, so has the need for computational methods capable of efficiently analyzing these increasingly large and information-rich datasets [2].

Machine learning (ML), particularly deep learning, has become a powerful tool for addressing this challenge. Modern ML workflows can automatically analyze entire WSIs, detect pathological features, and generate quantitative measurements difficult or impractical to obtain manually.

Traditionally, these approaches have relied on supervised learning, requiring large, carefully annotated datasets for each individual task. While highly successful, creating such datasets is often labor-intensive, expensive, and requires substantial annotation from a relatively small pool of domain experts, limiting the scalability of supervised approaches [3].

Recently, vision foundation models (FMs) have emerged as a promising approach for transferring pretrained visual representations to a wide range of image analysis tasks [4], [5]. Rather than training a new model for every application, FMs are pretrained on large and diverse image collections to learn general visual representations that can be transferred to a wide range of downstream tasks. One common approach is to keep the pretrained FM “frozen,” meaning its parameters are not modified, and use it to generate a numerical representation, or embedding, for each image. These embeddings capture visual features learned during pretraining and can be paired with known class labels to train a simple linear classifier, commonly referred to as a linear probe [6]. In computational pathology, this approach has shown impressive performance, especially in the cancer domain, across several applications while requiring substantially less task-specific training than developing a model end-to-end, raising the possibility of reducing the amount of annotated data required for downstream applications [4], [5].

Numerous vision FMs have recently been developed specifically for computational pathology, including UNI, Virchow/Virchow2, CONCH, Phikon, and Prov-GigaPath [6], [7], [8], [9], [10], [11].

Many of these models were pretrained predominantly on large collections of cancer and other clinical pathology images and have demonstrated strong performance across a variety of downstream pathology tasks. However, specialized disease domains, such as neurodegenerative disease pathologies, remain comparatively underrepresented in the pretraining and evaluation of current pathology FMs. This domain presents a distinct setting in which disease assessment relies on post-mortem brain tissue and a combination of hematoxylin and eosin (H&E), special histochemical, and immuno-histochemistry (IHC) stains to characterize disease-specific pathological hallmarks, including those used for Braak neurofibrillary tangle (NFT) staging, Thal amyloid phase, and CERAD scoring to pathologically diagnose Alzheimer disease [12], [13]. Consequently, it remains unclear how effectively representations learned largely from broader clinical and cancer pathology transfer to these specialized neurodegenerative disease tasks.

Recent independent benchmarking studies have also demonstrated pathology FMs’ generalization varies considerably across models and downstream tasks. Performance depends on the specific clinical task, adaptation strategy, and evaluation cohort, highlighting no single FM consistently outperforms others across all settings. While these studies have substantially advanced understanding of pathology FM generalization, they have focused primarily on common pathology tasks, particularly cancer-related applications, leaving transferability to specialized neurodegenerative disease pathology largely unexplored [14], [15], [16].

Domain-specific pretraining has also begun to address this gap. NeuroFM, for example, was trained specifically on datasets containing diverse neurodegenerative diseases and demonstrated improved performance over general-purpose pathology FMs on several neuropathology-specific tasks [17]. These findings suggest domain-specific pretraining may improve transfer to specialized neurodegenerative pathology; however, such models remain relatively recent and were not included in the present study.

Although linear probing is commonly used to evaluate frozen FM embeddings, alternative downstream strategies including k-nearest-neighbor classification and attention-based multiple-instance learning have also been applied in pathology datasets [18]. The present study focuses specifically on linear probing to evaluate the direct transferability of frozen representations, while more flexible downstream adaptation strategies are left for future work.

Direct comparisons between frozen foundation-model representations and conventional supervised convolutional neural networks (CNNs) remain limited as well, leaving an important question unanswered: under what conditions do pathology foundation models provide an advantage over traditional supervised learning?

To address these questions, we performed a systematic comparison of fourteen frozen feature extractors, including general-vision and pathology FMs, across four pathology image classification datasets (are there citations for the methods mentioned? If so, perhaps cite the foundational papers?). These included two specialized neurodegenerative disease immunohistochemistry datasets (with a focus on pathological hallmarks of Alzheimer disease: amyloid-β plaque classification and tau NFT pathology classification) and two hematoxylin and eosin (H&E) datasets, TIL (tumor infiltrating lymphocyte) and BACH (breast cancer histology)[19], [20]. Because both neurodegenerative disease datasets are IHC-based whereas both cancer-related datasets are H&E-based, pathology domain and staining modality are confounded in the present comparison and cannot be disentangled from one another. Frozen foundation-model linear probes were compared with a conventional supervised ResNet-50 CNN trained directly on image tiles. Finally, we investigated how CNN performance scaled with increasing amounts of labeled training data to better understand the relationship between supervised learning and frozen foundation-model representations. To facilitate reproducible evaluation of future foundation models and datasets, all experiments were implemented within a unified, dataset-agnostic evaluation pipeline that will be released as open source.

Together, these experiments evaluate the transferability of current pathology foundation models across distinct pathology domains and identify scenarios in which conventional supervised learning remains advantageous.

## 2. Materials & Methods

All experiments were performed using a unified evaluation pipeline designed to enable consistent comparison of multiple vision foundation models and a conventional supervised CNN across heterogeneous pathology datasets (Figure 1). Dataset-specific preprocessing was limited to conversion into a common dataset structure (i.e. metadata fields and allowable values, image format, etc.), after which all downstream processing, training, and evaluation followed an identical protocol (Figure 1). The code used during this work can be found freely available on GitHub (https://github.com/Gutman-Lab/biorxiv-vizcarra-2026). All experiments were conducted on a server running Ubuntu 24.04.4 LTS, with four NVIDIA L40S GPUs.

**Figure 1.**
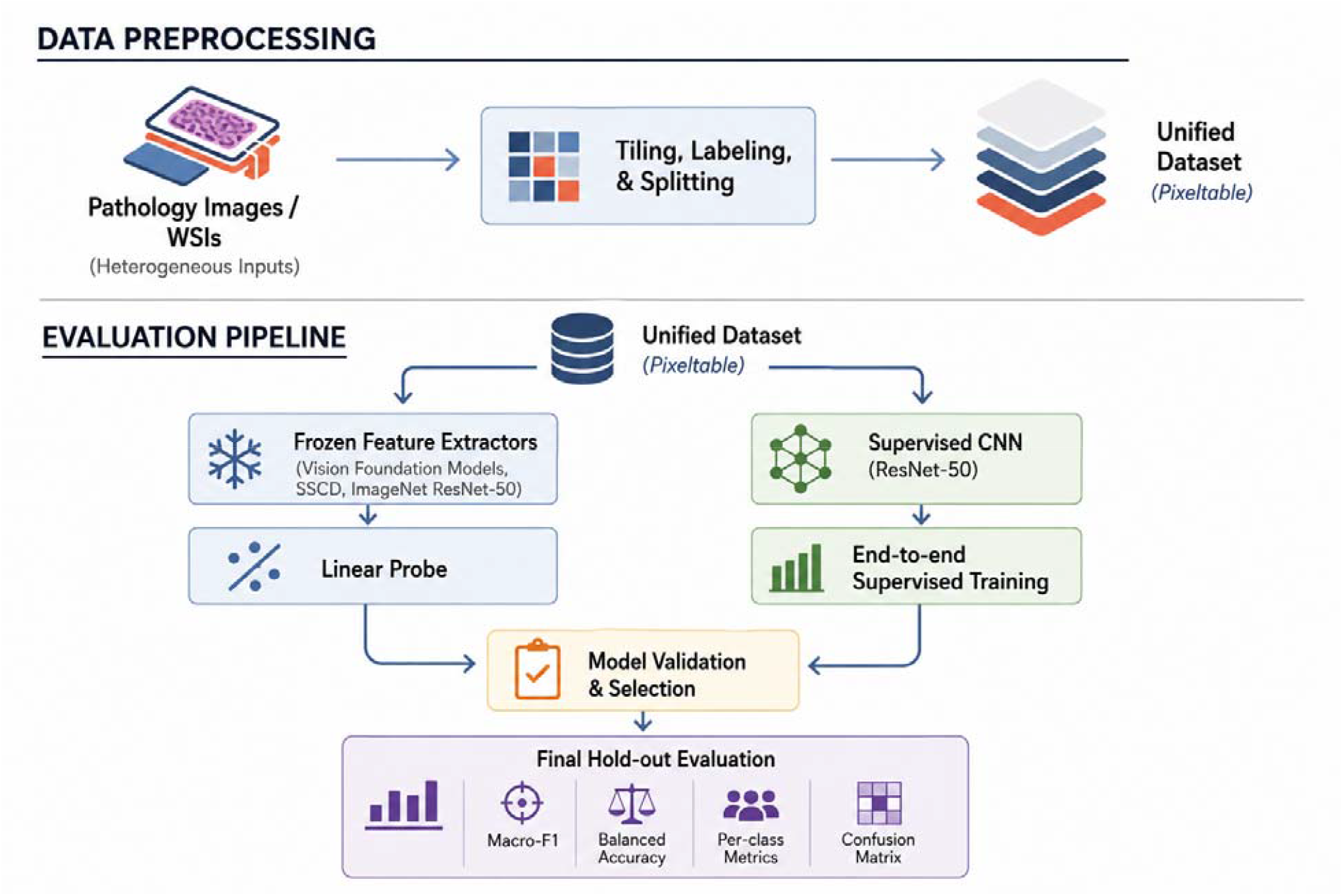
Overview of the experimental workflow used throughout this study. Dataset preprocessing converts each dataset into a common representation for downstream analysis within the Pixeltable database. Once loaded, all datasets are processed using an identical evaluation pipeline. Frozen vision foundation models are evaluated by generating image embeddings followed by linear-probe training (left path-blue boxes), while a supervised ResNet-50 CNN is trained end-to-end directly on image tiles (right path-green boxes). Both approaches use identical training, validation, and hold-out partitions, with model selection performed on the validation set and final performance reported on the hold-out test set. Figure artwork was generated with assistance from ChatGPT (OpenAI) and reviewed and edited by the authors.

### 2.1 Datasets

#### aBeta Dataset

The amyloid-β IHC dataset was obtained from Zenodo and was originally published by Tang *et al*. [21]. The WSIs used in that dataset were taken from the Alzheimer’s Disease Research Center (ADRC) at the University of California, Davis, from the temporal gyri of the brain.

Candidate pathologies were first identified using a custom blob-detection algorithm and presented to annotators for classification. Image regions were then extracted around the annotated objects to generate the image-level dataset. Each image could contain one or more labels corresponding to cored plaques, diffuse plaques, and/or cerebral amyloid angiopathy (CAA). Some images were considered negative because they have none of the pathologies present. To formulate a single-label classification task we excluded tiles that had more than one type of pathology available. We split the cohort by unique WSI into 60:20:20 splits (train:val:test). All images were scanned at 40x magnification using an Aperio AT2 and tiled at 256 by 256 pixels in RGB format. The original dataset was highly imbalanced with 80% diffuse, 16% negative, 2.5% cored, and 1% CAA (Table 1).

**Table 1.** Number of tiles in the amyloid-β dataset, including the tiles in each class for each split used during model development.

| *split* | *n_samples* | *CAA* | *Cored* | *Diffuse* | *Negative* |
| --- | --- | --- | --- | --- | --- |
| train | 30,998 | 392 | 770 | 24,935 | 4,901 |
| validation | 10,563 | 72 | 268 | 8,349 | 1,874 |
| hold-out | 11,181 | 118 | 256 | 9,094 | 1,713 |
| total | 52,742 | 582 | 1,294 | 42,378 | 8,488 |
| % | 100 | 1.1 | 2.5 | 80 | 16.1 |

#### Tau Dataset

The tau NFT IHC dataset was obtained from Emory University, and was previously published [22]. This dataset consisted of four brain regions: hippocampus, amygdala, temporal cortex, and occipital cortex. Images were scanned at 40x magnification using an Aperio scanner, selected regions of interest (ROIs) within the WSIs were exhaustively annotated for mature intraneuronal NFTs (iNFTs) and pre-NFTs. Annotators initially provided point annotations, which were subsequently converted semi-manually into bounding boxes. The annotated ROIs were then divided into non-overlapping 256 × 256 pixel tiles. We converted this dataset to a classification task by assigning labels to each tile based on presence or absence of each NFT type. To account for NFTs cut off when tiling, the label was ignored if the tile contained less than 50% of the original object in the ROI. This prevents assigning labels to tiles in which the object is only partially visible. Tiles with both types of NFTs were removed, those with neither type were marked as negative tiles and used as a class during training. The hold-out dataset was taken from the original dataset, and the rest were split into train and validation at a 80:20 ratio. This dataset was a fairly large dataset with nearly 250 thousand tiles in the dataset, with the majority of tiles being negative tiles and iNFT being substantially enriched compared to pre-NFT tiles (Table 2).

**Table 2.** The count of tiles in each dataset and the split across the classes for Tau dataset.

| *split* | *n_samples* | *pre-NFT* | *iNFT* | *negative* |
| --- | --- | --- | --- | --- |
| train | 193,766 | 5,666 | 20,178 | 167,922 |
| validation | 47,927 | 1,449 | 4,570 | 41,908 |
| hold-out | 6,134 | 50 | 455 | 5,629 |
| total | 247,827 | 7,165 | 25,203 | 215,459 |
| % | 100 | 2.9 | 10.2 | 87 |

#### Tumor Infiltrating Lymphocyte (TIL) Dataset

The Tumor Infiltrating Lymphocyte (TIL) dataset was obtained from Zenodo and was derived from H&E-stained diagnostic whole-slide images from The Cancer Genome Atlas (TCGA) spanning 23 cancer types [20]. The dataset is a curated subset of data generated by Abousamra et al. and Saltz et al. and contains 304,097 image patches [23], [24]. Because the TCGA images originated from multiple contributing institutions, the source WSIs were acquired using different slide scanners and staining protocols. Image patches are 100 by 100 pixels at 20X magnification, or approximately 50 by 50 µm field of view (0.5 µm/pixel resolution) [20].

Training labels were generated using a combination of pathologist-generated patch and region level annotations and model-generated annotations from a previously developed TIL classifier. For manual patch annotations, pathologists evaluated the center 50 by 50 µm region of larger 150 by 150 µm patches, with a patch considered TIL-positive when at least two lymphocytes or plasma cells were present. Pathologists also annotated TIL-positive and TIL-negative regions from which additional 50 by 50 µm patches were sampled. Model-generated annotations were obtained from classifications produced by the previously developed TIL model [23], [24].

The released dataset provides predefined training, validation, and hold-out patches, with TCGA participant identifiers used during partitioning to prevent images from the same participant from occurring in multiple splits. No stain normalization was applied to the released images. For this study, the cancer-specific cohorts were combined into a single dataset while preserving the original training, validation, and hold-out assignments. The final dataset contained 304,097 patches, of which approximately 18% were TIL-positive and 82% were TIL-negative (Table 3).

**Table 3.** The TIL dataset counts of each class across the three splits. This dataset is a binary dataset with two classes - positive or negative. Tiles from all cancers were combined into one large dataset for model development. Til + refers to images with two or more TILs present, TIL - refers to images without sufficient TIL presence.

| *split* | *n_samples* | *Til +* | *Til -* |
| --- | --- | --- | --- |
| train | 209,221 | 39,206 | 170,015 |
| validation | 38,601 | 5,203 | 33,398 |
| hold-out | 56,275 | 10,501 | 45,774 |
| total | 304,097 | 54,910 | 249,187 |
| % | 100 | 18.1 | 81.9 |

**Table 4.** BACH dataset, describing the counts of the dataset for each class and partition.

| *split* | *n_samples* | *Normal tissue* | *Benign lesion* | *In-situ carcinoma* | *Invasive carcinoma* |
| --- | --- | --- | --- | --- | --- |
| train | 240 | 60 | 60 | 60 | 60 |
| validation | 80 | 20 | 20 | 20 | 20 |
| hold-out | 80 | 20 | 20 | 20 | 20 |
| total | 400 | 100 | 100 | 100 | 100 |
| % | 100 | 25 | 25 | 25 | 25 |

#### BACH Dataset

Finally, we included the well-established BACH (BreAst Cancer Histology) benchmark introduced as part of the ICIAR (International Conference on Image Analysis and Recognition) 2018 Grand Challenge. This dataset consists of H&E-stained breast tissue images categorized into four classes: normal tissue, benign lesions, *in-situ* carcinoma, and invasive carcinoma. A total of 400 images are provided, with 100 images per class. The dataset is distributed as fixed-size microscopy images and therefore required no additional tiling for this study. For this study, images were randomly divided into training, validation, and hold-out sets using a stratified 60:20:20 split to preserve equal class representation across all partitions [19], [25].

For a comparison of the datasets see Table 5.

**Table 5.**
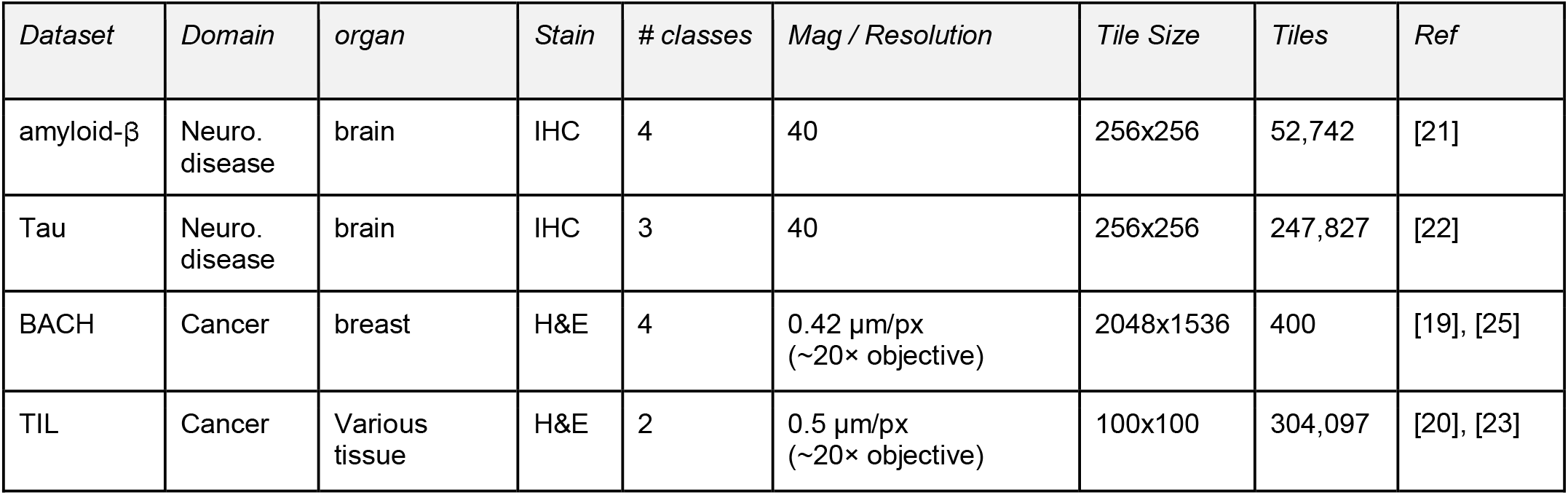
Comparison of the datasets used in this project.

| *Dataset* | *Domain* | *organ* | *Stain* | *# classes* | *Mag / Resolution* | *Tile Size* | *Tiles* | *Ref* |
| --- | --- | --- | --- | --- | --- | --- | --- | --- |
| amyloid-β | Neuro. disease | brain | IHC | 4 | 40 | 256x256 | 52,742 | [21] |
| Tau | Neuro. disease | brain | IHC | 3 | 40 | 256x256 | 247,827 | [22] |
| BACH | Cancer | breast | H&E | 4 | 0.42 $\mu\text{m}/\text{px}$<br>($\sim 20\times$ objective) | 2048x1536 | 400 | [19], [25] |
| TIL | Cancer | Various tissue | H&E | 2 | 0.5 $\mu\text{m}/\text{px}$<br>($\sim 20\times$ objective) | 100x100 | 304,097 | [20], [23] |

### 2.2 Experimental Pipeline

To ensure a fair comparison across datasets and models, all experiments were performed using a unified evaluation pipeline (Figure 1). Because each dataset originated from different sources and annotation formats, a small amount of dataset-specific preprocessing was required to convert each dataset into a common representation. This preprocessing included operations such as tiling WSIs, converting annotations into classification labels, and constructing standardized train, validation, and hold-out splits [19], [20], [21], [22], [23], [25]. No stain normalization was applied during dataset preprocessing or model evaluation [26]. Beyond the dataset-specific preprocessing steps described above, images were otherwise used in their original form. Once processed, all datasets were stored in a unified local database implemented using Pixeltable (all code is available on GitHub: https://github.com/Gutman-Lab/pixelEmbed).

After preprocessing, all datasets followed an identical downstream evaluation workflow. In the first branch, frozen image embeddings were generated using each vision foundation model (see table 6 for each model) and used to train a linear probe for downstream classification. In the second branch, a supervised ResNet-50 CNN was trained end-to-end directly on the image tiles. Both approaches used identical training, validation, and hold-out partitions, ensuring all comparisons were performed under the same experimental conditions.

**Table 6.** Information on the foundation models and frozen weights used for the linear probes.

| *Model* | *Model Type* | *Embedding Dimension* | *# of Params* | *Pretraining Data* | *Ref* |
| --- | --- | --- | --- | --- | --- |
| CLIP ViT-B/32 | General Vision FM | 512 | 87.8 M | Internet image-text pairs | [27] |
| CLIP ViT-B/16 | General Vision FM | 512 | 86.2 M | Internet image-text pairs | [27] |
| CLIP ViT-L/14 | General Vision FM | 768 | 304 M | Internet image-text pairs | [27] |
| DINOv2-B | General Vision FM | 768 | 86.6 M | General natural images | [28] |
| DINOv2-L | General Vision FM | 1024 | 304.4 M | General natural images | [28] |
| CONCH | Pathology FM | 512 | 89.9 M | 1.17M pathology image-caption pairs | [11] |
| UNI | Pathology FM | 1024 | 303.3 M | H&E WSIs | [10] |
| UNI2-h | Pathology FM | 1536 | 681.4 M | H&E + IHC WSIs | [10] |
| Phikon-v2 | Pathology FM | 1024 | 303.4 M | 60k H&E WSIs | [8] |
| Virchow | Pathology FM | 2560 | 631.2 M | Histopathology H&E WSIs | [6] |
| Virchow2 | Pathology FM | 2560 | 631.2 M | Histopathology H&E & IHC WSIs | [9] |
| Prov-GigaPath | Pathology FM | 1536 | 1.13 B | Histopathology H&E & IHC WSIs | [7] |
| ResNet50 (Frozen) | ImageNet CNN | 2048 | 23.5 M | ImageNet | [29], [30] |
| SSCD | Self-supervised CNN | 1024 | 44.2 M | General natural images | [31] |

### 2.3 Foundation Models

Most models were obtained from the Hugging Face (HF) Hub. Because model weights were distributed through different software frameworks—including Hugging Face Transformers, timm/HF Hub, the Mahmood Lab CONCH/OpenCLIP package, and a local TorchScript checkpoint for SSCD—we instantiated each model using its corresponding official implementation rather than a single standardized API.

Pathology foundation models were UNI (HF/Transformers); UNI2-h, Virchow, Virchow2, and Prov-GigaPath (HF weights via timm); CONCH (HF weights via Mahmood Lab’s CONCH/OpenCLIP package); and Phikon-v2 (HF/Transformers) [6], [7], [8], [9], [10], [11], [27]. General-purpose foundation models were OpenAI CLIP (HF/Transformers via Pixeltable) and Meta DINOv2 (HF/Transformers) [28].

Two additional frozen encoders served as non-foundation-model baselines for the linear-probe experiments: ImageNet-pretrained ResNet-50 (timm) and Meta SSCD, a self-supervised copy-detection model that extracts image-similarity features from natural images (local TorchScript checkpoint) [29], [30], [31]. Model details are given in Table 6.

### 2.4 Linear Probe

With the backbone frozen, each tile is represented by a fixed embedding that is L2-normalized and then classified by a single linear layer mapping embedding dimension to the number of classes. The probe is optimized using stochastic gradient descent with class-weighted cross-entropy loss. Balanced class weights are computed from the training split, and a fixed batch size of 256 tiles is used throughout all experiments. Hyperparameters, including the learning rate, are selected by a 15-epoch grid search using validation macro-F1. The selected configuration is then retrained from scratch for up to 100 epochs with early stopping. Early-stopping parameters (patience = 10 epochs; minimum improvement in validation macro-F1 = 0.005) were fixed a priori and were not included in hyperparameter optimization. Early stopping was used to reduce unnecessary computation after validation performance plateaued. The best validation checkpoint was evaluated once on the hold-out test set.

### 2.5 CNN Baseline

The CNN path fine-tunes a ResNet-50 initialized from ImageNet-pretrained weights end-to-end on the same train/validation/hold-out tile splits used for the linear probes. Hyperparameters are selected by a grid search of 18 configurations—three learning rates, two weight decays, and three class-weighting schemes (unweighted, square-root inverse-frequency, and effective-number)—each trained for 15 epochs; the setting with the highest validation macro-F1 is retained. That configuration is then retrained from scratch for up to 100 epochs with the same early stopping strategy used for the linear probe. Like the linear probe, the best validation checkpoint was evaluated once on the hold-out test set.

Despite training the CNN end-to-end, the computational requirements remained relatively modest. For the largest CNN training experiment, using the complete Tau training dataset of 193,766 tiles, the final selected model required approximately 7 hours to train, while evaluation of the full 18-configuration hyperparameter grid required approximately 31 hours on a single NVIDIA L40S GPU.

### 2.6 Evaluation

Each trained model was evaluated once on the hold-out test set. Performance was quantified using macro-F1 score, per-class precision, recall, F1 score, and confusion matrices. For frozen feature extractors, computational efficiency was assessed using mean embedding-generation time per image. For each model and dataset, total embedding-generation wall-clock time was divided by the number of images processed. Timing was performed on the same computing infrastructure used for the other experiments, and values are reported in milliseconds per tile (Table 7). Macro-F1 served as the primary evaluation metric and was also used for hyperparameter selection because three of the four datasets exhibited substantial class imbalance. Unlike overall accuracy, macro-F1 weights each class equally regardless of class frequency, making it a more appropriate metric for evaluating performance on imbalanced classification tasks.

**Table 7.** Average embedding time for each FM (plus the frozen ResNet50 and SSCD model) across the four datasets tested. All values are measured in milliseconds per tile.

| *Model* | *amyloid-β* | *Tau* | *BACH* | *TIL* |
| --- | --- | --- | --- | --- |
| CLIP ViT-B/32 | 6.19 | 4.85 | 23.67 | 3.66 |
| CLIP ViT-B/16 | 7.47 | 5.75 | 23.11 | 4.70 |
| CLIP ViT-L/14 | 12.26 | 10.30 | 29.43 | 9.32 |
| CONCH | 7.90 | 6.67 | 45.06 | 5.08 |
| UNI | 10.80 | 8.93 | 31.05 | 7.72 |
| UNI2-h | 19.42 | 17.27 | 37.30 | 16.29 |
| DINOv2-L | 10.59 | 9.21 | 25.88 | 8.67 |
| DINOv2-B | 6.54 | 5.28 | 23.93 | 4.61 |
| Phikon-v2 | 10.62 | 8.53 | 26.76 | 7.62 |
| ResNet50 (Frozen) | 7.01 | 5.16 | 37.20 | 4.19 |
| Virchow | 19.72 | 17.08 | 45.33 | 16.09 |
| Virchow2 | 19.91 | 17.30 | 45.08 | 16.19 |
| Prov-GigaPath | 23.73 | 21.09 | 48.93 | 19.98 |
| SSCD | 7.03 | 5.28 | 27.06 | 4.18 |

### 2.7 Scaling Experiment

To investigate the relationship between training dataset size and model performance, we performed a scaling experiment using the Tau NFT dataset, which contained 193,766 training tiles. Hyperparameters selected during the full-dataset experiments were fixed throughout the scaling analysis. Nested subsets containing 1%, 5%, 10%, 25%, 50%, and 100% of the original training data were generated, with each larger subset containing all samples from the previous subset. Subsets were drawn by seeded random sampling without class stratification; consequently, class proportions could vary across subsets. In practice, train-set class frequencies remained similar to the full training distribution (Table 8). Frozen linear probes and the supervised CNN were trained on identical image subsets to ensure a fair comparison between approaches. Validation and hold-out partitions remained unchanged, and all models were evaluated using the same protocol described above.

**Table 8.** Scaling experiment dataset subset information. For each subset we report the total number of images in the training dataset, number of images in each class, and the percentage of images of each class in the subset.

| % Train | Total Images | Negative | Negative % | Pre-NFT | Pre-NFT % | iNFT | iNFT % |
| --- | --- | --- | --- | --- | --- | --- | --- |
| 1 | 1,938 | 1,688 | 87.1 | 70 | 3.6 | 180 | 9.3 |
| 5 | 9,688 | 8,385 | 86.6 | 282 | 2.9 | 1,021 | 10.5 |
| 10 | 19,376 | 16,799 | 86.7 | 560 | 2.9 | 2,017 | 10.4 |
| 25 | 48,440 | 41,906 | 86.5 | 1,382 | 2.9 | 5,152 | 10.6 |
| 50 | 96,880 | 83,941 | 86.6 | 2,771 | 2.9 | 10,168 | 10.5 |
| 100 | 193,766 | 167,922 | 86.7 | 5,666 | 2.9 | 20,178 | 10.4 |

## 3. Results

### 3.1 Overall performance across pathology datasets

Across the four datasets, macro-F1 and balanced accuracy showed broadly similar performance patterns among the evaluated models (Table 9 & Supplementary Figure 1). On both IHC datasets, the supervised ResNet-50 CNN achieved the highest macro-F1 score. For the amyloid-β dataset, the CNN achieved a macro-F1 of 0.762. The highest-performing frozen feature extractor was SSCD with a macro-F1 of 0.604, while CONCH was the highest-performing pathology foundation model at 0.593. Similarly, on the Tau NFT dataset, the CNN achieved a macro-F1 of 0.758, compared with 0.580 for the best-performing frozen foundation-model linear probe, Prov-GigaPath.

**Table 9.**
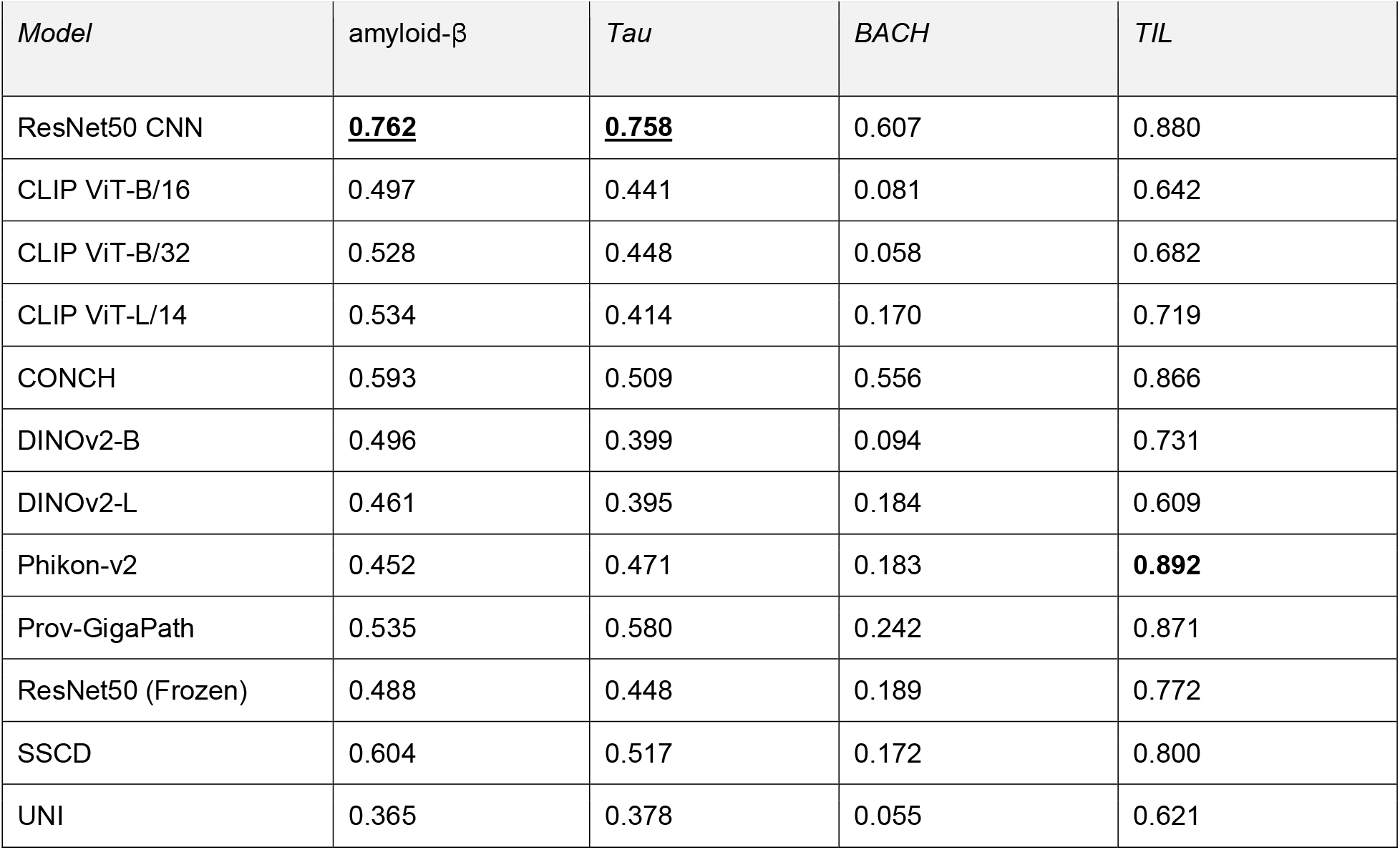
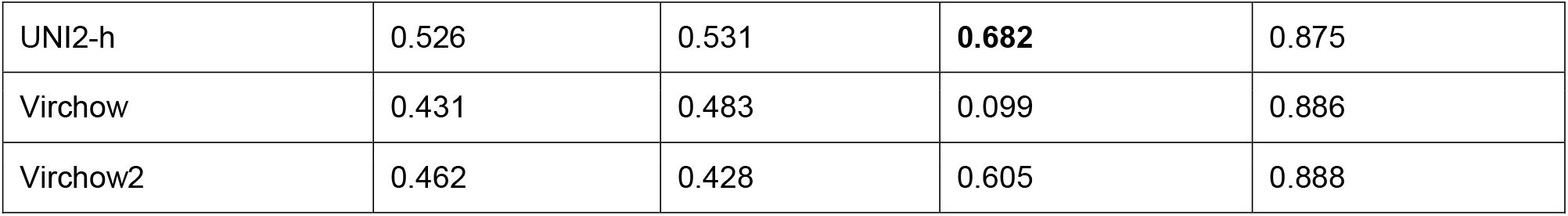
Macro F1 scores for each model and dataset, including all frozen FM, frozen non-FMs, and the ResNet50 trained end-to-end. The macro F1 score for each dataset is bolded, and underlined if it is also the ResNet50 CNN.

A different performance pattern was observed for the cancer based H&E datasets. On BACH, UNI2-h achieved the highest macro-F1 score of 0.682, followed by the supervised CNN at 0.607 and Virchow2 at 0.605. On the larger TIL dataset, several pathology foundation models achieved performance comparable to or slightly exceeding the CNN. Phikon-v2 achieved the highest macro-F1 at 0.892, followed by Virchow2 at 0.888 and Virchow at 0.886, while the CNN achieved 0.880. Thus, the relative performance of frozen representations and supervised CNN training differed substantially across pathology domains: the CNN showed a clear advantage on both neurodegenerative disease IHC tasks, whereas pathology FMs were competitive with or superior to the CNN on the two cancer based H&E datasets (Table 9. Per-class F1 scores for all models are provided in the Supplementary Figures.)

Given the substantial class imbalance in the amyloid-β dataset, we additionally examined per-class F1 scores for the supervised CNN and CONCH, the highest-performing pathology FM linear probe (Table 10). The CNN achieved higher F1 scores across all four classes, with the largest difference observed for the relatively rare cored-plaque class (0.669 versus 0.252). Performance was also higher for CAA (0.556 versus 0.410), indicating the overall CNN advantage was not driven solely by performance on the dominant diffuse-plaque class.

**Table 10.** Per-class F1 for each class of the amyloid-β dataset, comparing the results from the ResNet50 CNN trained end-to-end and the best performing FM linear probe.

| *Model* | *Negative* | *Cored* | *Diffuse* | *CAA* |
| --- | --- | --- | --- | --- |
| ResNet50 CNN | 0.854 | 0.669 | 0.967 | 0.556 |
| CONCH | 0.777 | 0.252 | 0.934 | 0.410 |

Embedding-generation time varied substantially among frozen feature extractors (Table 7). These differences were particularly relevant when models achieved similar predictive performance. On the TIL dataset, Phikon-v2 achieved the highest macro-F1 (0.892) with a mean embedding time of 7.62 ms per tile, compared with 16.19 ms for Virchow2 and 16.09 ms for Virchow, which achieved macro-F1 scores of 0.888 and 0.886, respectively. Thus, computational efficiency may provide an additional practical consideration when selecting among frozen feature extractors with similar downstream performance.

### 3.2 Scaling experiment

To investigate whether the performance gap between frozen foundation-model representations and supervised CNNs could be explained by training dataset size, we performed a scaling experiment using the Tau NFT dataset. The supervised CNN consistently outperformed the best frozen linear probes across all training set sizes (Figure 3). Using only 1% of the available training data, the CNN achieved a macro-F1 score of 0.620, exceeding the best frozen foundation-model linear probe trained using the complete training dataset (Prov-GigaPath, macro-F1 = 0.580).

**Figure 3.**
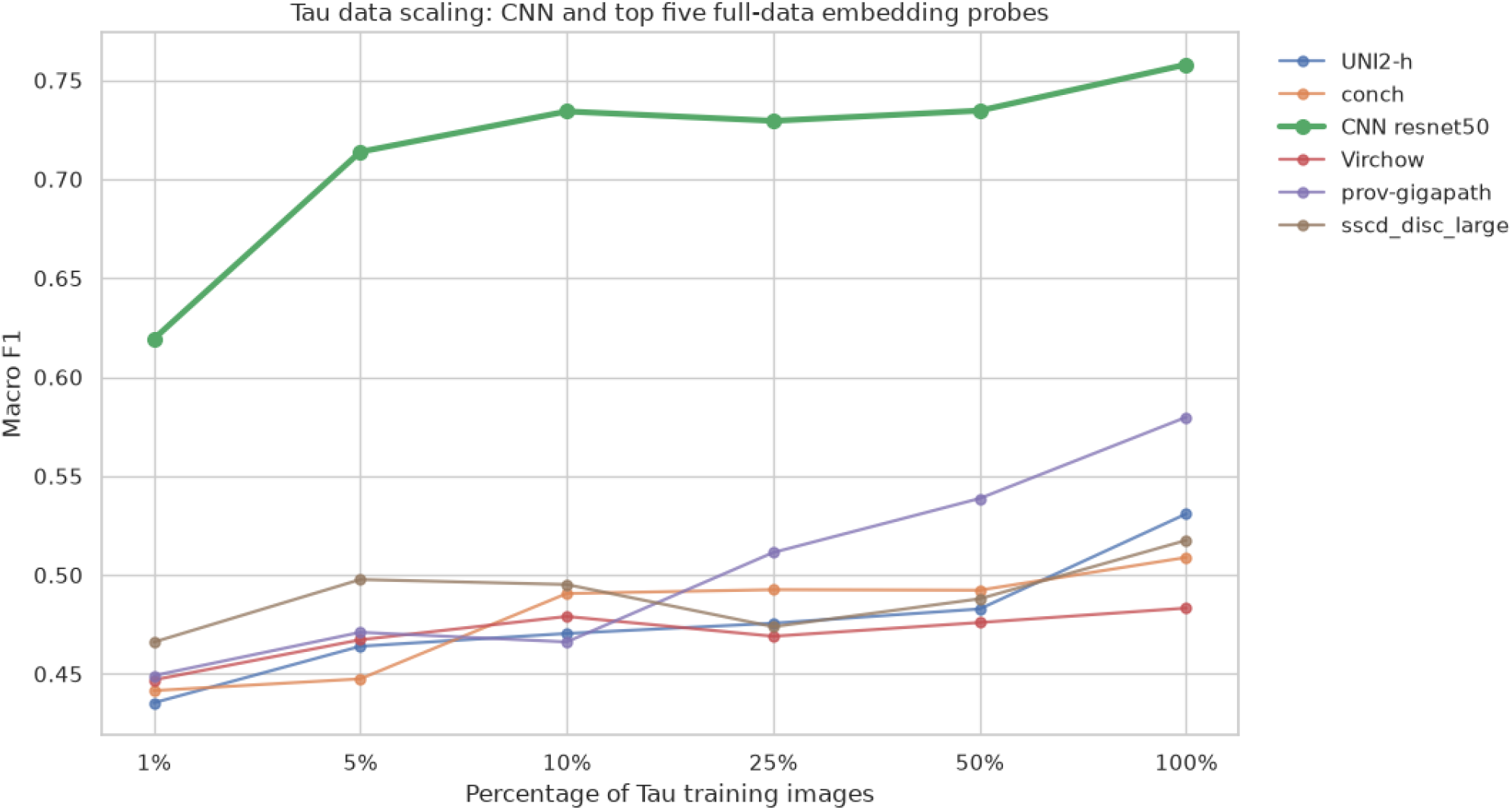
Scaling experiment results showing the macro F1 score on the holdout set of the tau NFT dataset. The top five performing linear probes and the CNN are shown across the different values of percentage of data retained for training.

Several pathology foundation models exhibited gradual improvements in macro-F1 as additional training data became available. For example, Prov-GigaPath improved from approximately 0.45 at 1% of the training data to 0.58 when trained on the full dataset. In contrast, several other models demonstrated only modest performance gains despite a 100-fold increase in the amount of labeled training data (Supplementary Fig 5). Across all training set sizes, the supervised CNN maintained a substantial performance advantage over the frozen linear probes.

## 4. Discussion

### 4.1 Domain-dependent transferability of pathology foundation models

The results of this study demonstrate the transferability of frozen pathology FMs varies considerably across evaluated datasets. On both cancer based H&E datasets (BACH and TIL), pathology FMs consistently achieved performance comparable to, and in some cases exceeding, the supervised ResNet-50 CNN. Although the two datasets represent different classification tasks—multiclass tissue classification in BACH and binary tumor infiltrating lymphocyte detection in TIL—the overall performance pattern was similar, with pathology FMs occupying the top-performing positions.

In contrast, the same pattern was not observed on the two neurodegenerative disease datasets. For both the amyloid-β and Tau classification tasks, the supervised CNN substantially outperformed all frozen FM linear probes. Notably, the CNN advantage on the amyloid-β dataset extended across both common and rare classes, with the largest improvement observed for cored plaques. This performance gap remained consistent despite the large size o the Tau dataset, suggesting the observed difference cannot be attributed solely to limited training data.

Another notable observation was the relative performance of pathology-specific and general vision FMs. On the H&E cancer datasets, pathology FMs consistently outperformed general vision models such as CLIP and DINOv2. However, this distinction was much less pronounced on the neurodegenerative disease datasets, where both groups of frozen foundation models achieved similar performance despite their different pretraining domains.

More broadly, these findings are consistent with growing evidence that pathology foundation models inherit biases from their pretraining data composition rather than learning universally transferable representations. For example, independent benchmarking of Prov-GigaPath has shown particularly strong performance on lung pathology, consistent with the fact that approximately 45% of its pretraining data originated from lung cancer specimens. These observations suggest downstream performance is influenced not only by stain diversity but also by the representation of specific tissues and disease processes during pretraining [16].

Taken together, these results suggest the transferability of frozen pathology foundation models depends strongly on the relationship between the downstream task and the data used during pretraining. While pathology FMs transferred effectively to the H&E cancer datasets evaluated here, this advantage was not observed for the specialized IHC tasks. Differences in annotation and sampling strategies across datasets may also have influenced model performance; however, these factors are confounded with the individual datasets and cannot be isolated in the present study.

### 4.2 Why do pathology foundation models perform differently across domains?

One possible explanation for the observed performance differences is the composition of the datasets used to pretrain current pathology FMs. Several widely used models were trained predominantly or exclusively on H&E histopathology, and primarily or at least highly enriched for cancer-related cohorts. UNI, for example, was pretrained on more than 100 million image patches from over 100,000 diagnostic H&E WSIs spanning 20 tissue types, while Virchow was trained on approximately 1.5 million H&E WSIs from Memorial Sloan Kettering Cancer Center. Phikon-v2 was similarly trained on 460 million pathology tiles collected from more than 100 publicly available cohorts covering over 30 cancer sites [6], [8], [10].

More recent FMs have incorporated broader staining diversity. UNI2-h was pretrained on more than 200 million tiles from over 350,000 H&E and IHC slides (taken from HuggingFace’s checkpoint provided by Mahmood lab), while Prov-GigaPath was trained on approximately 1.3 billion tiles from 171,189 H&E and IHC pathology slides across 31 tissue types [7], [10].

Virchow2 likewise expanded pretraining to 3.1 million WSIs spanning multiple stains and nearly 200 tissue types [9]. CONCH is notable for using more than 1.17 million pathology image-caption pairs, including substantial numbers of IHC and special-stain images in addition to H&E [11].

Despite this increasing diversity, the extent to which specialized neurodegenerative morphology and IHC pathology are represented in these pretraining datasets is generally unclear. The inclusion of brain tissue or IHC slides does not necessarily imply substantial exposure to structures such as amyloid-β plaques, CAA, pre-NFTs, and/or mature NFTs. These highly specific pathological objects may therefore require representations that differ from those learned predominantly from broader histopathology and cancer datasets. This may help explain why pathology-specific FMs showed a clear advantage over general vision FMs on the H&E datasets but not on the neurodegenerative disease IHC tasks evaluated here.

These findings suggest simply increasing the diversity of pathology FM pretraining data may not be sufficient; representation of specialized disease domains and staining patterns may also be important for downstream transfer. Larger and more diverse training collections could therefore improve FM performance on targeted neurodegenerative pathology tasks.

### 4.3 Error Analysis

To better understand the performance difference observed on the Tau dataset, we compared the confusion matrices of the supervised CNN and the best-performing pathology FM (Prov-GigaPath) (Figure 4). The largest difference between the two models was not in the classification of mature NFTs, where both achieved similar performance, but rather in their handling of negative tissue and early pathology. The supervised CNN correctly classified 97% of negative tiles compared with 89% for Prov-GigaPath, indicating substantially fewer false-positive predictions. In addition, the FM more frequently confused pre-NFTs with mature NFTs (24% versus 10% for the CNN), suggesting subtle distinctions between early and mature pathology are less well represented in the frozen embedding space. Together, these observations suggest, for this Tau dataset and Prov-GigaPath comparison, part of the performance gap may reflect the CNN’s improved discrimination of subtle pathological features and normal tissue rather than improved recognition of mature NFTs themselves.

**Figure 4.**
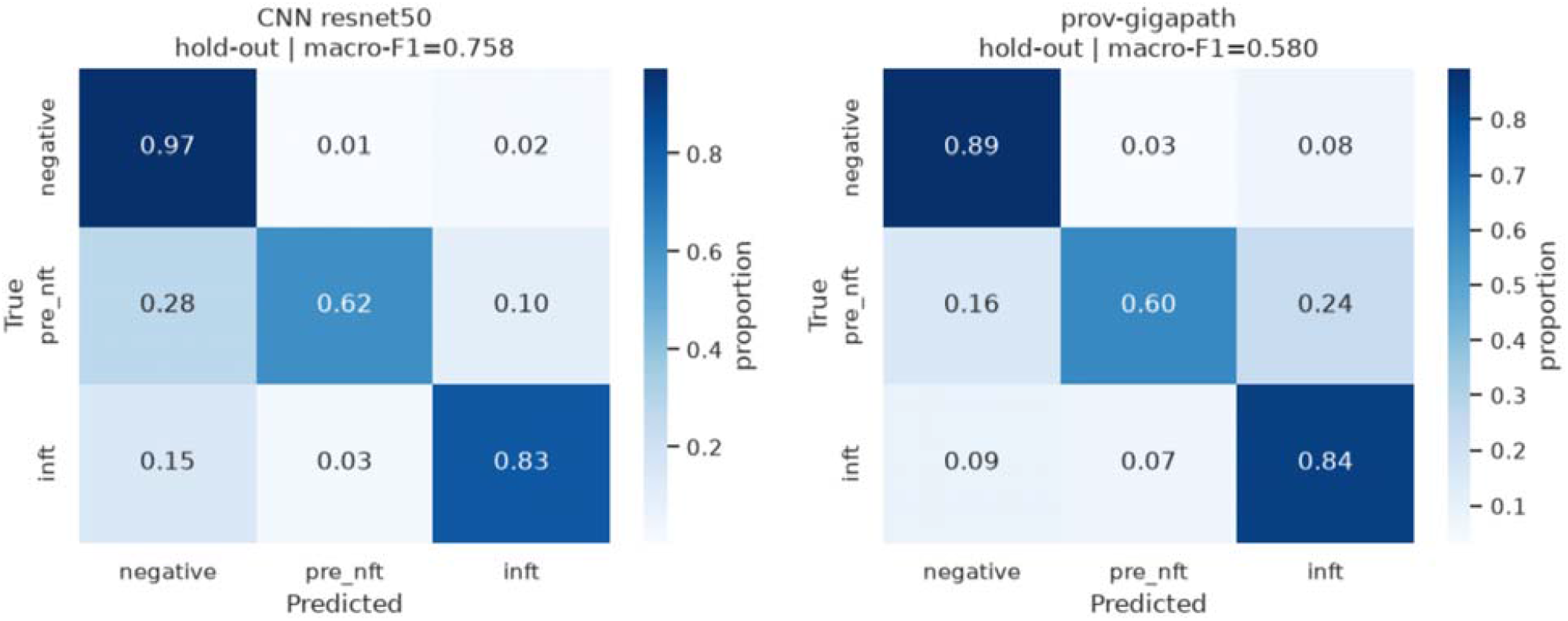
Row-normalized confusion matrices for the Tau hold-out test set comparing the supervised ResNet-50 CNN with the best-performing pathology FM, Prov-GigaPath. Values represent the proportion of predictions within each true class.

### 4.4 What does the scaling experiment tell us?

One of the primary advantages of FMs is they can be applied to a wide variety of downstream tasks without retraining the feature extractor, substantially reducing the computational cost and annotation effort typically required for supervised learning. Because our study relied exclusively on publicly available datasets, an important question was whether the superior performance of the supervised CNN simply reflected the availability of sufficiently large training datasets.

The scaling experiment on the Tau dataset suggests this is unlikely to be the sole explanation. Despite using only 1% (1,938 tiles) of the available training data, the supervised CNN still outperformed the best frozen FM trained using the complete dataset. This observation indicates the performance gap cannot be explained solely by the quantity of labeled training data available to the downstream classifier.

Increasing the amount of training data did improve the performance of some FMs. For example, Prov-GigaPath improved by approximately 15 percentage points in macro-F1 between the 1% and 100% training subsets. However, several models, particularly the general vision FMs, showed relatively little improvement as additional training data became available (Figure 3 and Supplementary Figure 5). This suggests some frozen representations contain features that become increasingly useful with additional supervision, whereas others may not encode sufficient task-relevant information for this specialized neurodegenerative disease classification task.

Together, these findings support the hypothesis that the observed performance differences are influenced not only by the amount of downstream training data but also by the quality and relevance of the representations learned during FM pretraining. Whether this reflects limited representation of organ and disease specific morphology, staining differences, ways in which items were annotated, or other characteristics of the pretraining datasets are important questions for future work.

### 4.5 Practical implications

The results of this study suggest different strategies depending on the pathology domain of interest. For common H&E pathology tasks, frozen pathology foundation models provide an excellent starting point and may achieve performance comparable to, or exceeding, conventionally trained CNNs. Consequently, when beginning a new H&E classification project, evaluating frozen FM embeddings with a linear probe may provide a rapid baseline before committing resources to the collection of large, task-specific annotated datasets.

In contrast, our results indicate the frozen FM paradigm evaluated in this study is less effective for the specialized classification tasks examined here. For studies requiring high-performance detection of neurodegenerative pathology, collecting a well-annotated dataset and training a supervised model end-to-end remains a strong approach. Such datasets may also prove valuable for future pretraining or fine-tuning of neuropathology-specific foundation models capable of improved transfer to these specialized tasks.

Importantly, both approaches remained computationally practical using standard GPU infrastructure. While downstream linear probes could typically be trained within hours, end-to-end CNN training was also relatively modest in computational cost. Even for the largest dataset evaluated, the final CNN required approximately 7 hours of training, with the complete hyperparameter search requiring approximately 31 hours on a single NVIDIA L40S GPU. Thus, conventional supervised CNN training remains a practical alternative to frozen FM approaches when sufficiently annotated data are available and should not necessarily be excluded on the basis of computational cost alone.

Beyond the benchmarking results themselves, this work provides a unified and standardized evaluation pipeline for comparing vision foundation models and supervised CNNs across heterogeneous pathology datasets. Because only minimal dataset-specific preprocessing is required before insertion into the common pipeline, new datasets and future foundation models can be incorporated with relatively little effort. Although this study focused on frozen linear probes and supervised CNN classification, the framework was designed to be extensible and already supports future additions, including evaluation of alternative CNN architectures, multi-label classification tasks, nonlinear probes, and transformer-based classifiers. Importantly, training the downstream linear probes required relatively modest computational resources and could typically be completed within hours rather than the weeks or months often associated with training large models. This makes evaluation of pretrained FMs accessible to laboratories with standard GPU computing infrastructure, without requiring the computational resources needed to train large models from scratch. By releasing the complete pipeline as open source, we hope to facilitate standardized benchmarking and simplify future comparisons as new pathology foundation models become available.

### 4.6 Limitations

Several limitations should be considered when interpreting the results of this study. First, our comparison focused exclusively on frozen foundation-model embeddings evaluated using linear probes. We did not investigate end-to-end fine-tuning of the foundation models, nonlinear probe architectures, or more extensive optimization of the linear-probe training procedure.

Consequently, the observed performance differences should be interpreted as comparisons between frozen foundation-model representations and a conventionally trained supervised CNN, rather than the maximum achievable performance of each foundation model. However, this comparison also reflects one of the fundamental questions motivating this study. A FM trained or adapted specifically for neuropathology/neurodegenerative diseases might reasonably be expected to outperform a relatively simple supervised CNN. Here, however, we sought to determine whether existing pretrained foundation models, when used without task-specific adaptation, provide a meaningful advantage over training a conventional classifier directly on the available labeled data, even when relatively few training samples are available.

Second, although four task specific datasets spanning both cancer and neurodegenerative tasks were evaluated, they represent only a subset of the many publicly available pathology datasets. Likewise, only a single CNN architecture (ResNet-50) was evaluated, and all datasets were formulated as single-label classification tasks, including conversion of the originally multi-label amyloid-β dataset into a single-label classification problem. These design choices were made to provide a consistent and standardized benchmark across all datasets and models.

In addition, all models were evaluated using a single train/validation/hold-out split and a single training run, so the reported results do not capture variability arising from model initialization, data sampling, or the specific dataset partition. This limitation is particularly relevant for small datasets such as BACH and for comparisons in which performance differences are modest.

Pathology domain and staining modality are also confounded in the present study: both neurodegenerative disease datasets use immunohistochemistry (IHC), whereas both cancer datasets use H&E staining. The datasets additionally differ in annotation procedures, tissue sampling, organ type, and other methodological characteristics. Because these factors co-vary across a limited number of datasets, their individual contributions to the observed performance differences cannot be determined independently. Although the performance patterns were consistent across the two datasets within each group, the present study cannot establish whether these differences arise primarily from pathology domain, organ type, sampling strategy, staining modality, annotation method, or interactions among these factors. Future work incorporating additional datasets spanning different disease domains and staining modalities, ideally with more comparable sampling and annotation procedures, will help disentangle these effects.

### 4.7 Future work

Several directions will be explored to further understand the transferability of pathology FMs. First, we plan to expand the benchmark by incorporating additional datasets, with particular emphasis on neurodegenerative disease H&E datasets and cancer IHC datasets. This will allow the effects of pathology domain and staining modality to be investigated independently, addressing one of the primary limitations of the current study.

Second, we will extend the scaling analysis to multiple datasets. Rather than evaluating only percentages of the available training data, future experiments will begin with a fixed number of annotated images and progressively increase the training set size. This will provide a more direct comparison of the data efficiency of supervised CNNs and frozen FM representations across different pathology tasks.

Another important direction is the evaluation of recently developed neuropathology-specific FMs. These models provide an opportunity to investigate whether domain-specific pretraining improves transferability across related neurodegenerative pathology tasks. For example, it will be interesting to determine whether an FM pretrained on amyloid-β and tau pathology transfers effectively to other neurodegenerative proteinopathies involving TDP-43 or α-synuclein, or whether end-to-end supervised training continues to provide superior performance [32].

Finally, future work will investigate alternative downstream adaptation strategies, including nonlinear probes, transformer-based classifiers, and end-to-end fine-tuning of foundation models. The evaluation pipeline developed in this work will also be extended beyond single-label classification to additional pathology tasks, including multi-label classification, object detection, and semantic segmentation.

## 5. Conclusions

This study systematically evaluated fourteen frozen feature extractors using frozen linear probes and compared their performance against a conventional supervised ResNet-50 CNN across four pathology classification datasets spanning both cancer focused H&E and neurodegenerative disease immunohistochemistry (IHC) tasks. While pathology foundation models achieved performance comparable to, and in some cases exceeding, the supervised CNN on the H&E datasets, the CNN consistently outperformed all frozen foundation-model representations on the two neurodegenerative disease IHC datasets. Furthermore, the scaling experiment demonstrated performance differences could not be explained solely by the availability of large annotated training datasets, as the CNN trained with only 1% of the Tau training data still exceeded the best frozen foundation model trained on the complete dataset.

Together, these findings suggest the transferability of frozen representations from current pathology FMs is strongly dependent on the downstream pathology domain and may be insufficient for specialized disease-specific classification tasks. In addition to these benchmarking results, this work provides an open, standardized evaluation pipeline that enables standardized comparison of vision foundation models and supervised CNNs across heterogeneous pathology datasets. We hope this framework will facilitate future benchmarking efforts and support continued development of pathology foundation models as new datasets, architectures, and adaptation strategies become available.

## Acknowledgements

ChatGPT (OpenAI) was used during development of the software pipeline to assist with code generation, debugging, performance optimization, and workflow orchestration. ChatGPT was also used during preparation of the manuscript to assist with editing, organization, and refinement of author-written scientific content. All generated code, analyses, interpretations, and manuscript text were reviewed and validated by the authors, who take full responsibility for the final work.

Research reported in this publication was supported by the National Institute of Neurological Disorders and Stroke of the National Institutes of Health under award number U24NS133949 and the National Institute on Aging under award number R01AG062517, in addition to support from the Chan Zuckerberg Initiative DAF [2024□351073] (B.N.D., D.A.G., C.C.), an advised fund of the Silicon Valley Community Foundation.*The content is solely the responsibility of the authors and does not necessarily represent the official views of the National Institutes of Health*.

