## Supplemental File for "Evaluating the Transferability of Pathology Foundation Models Across Cancer-related H&E Neurodegeneration-related Immunohistochemical Classification Tasks"

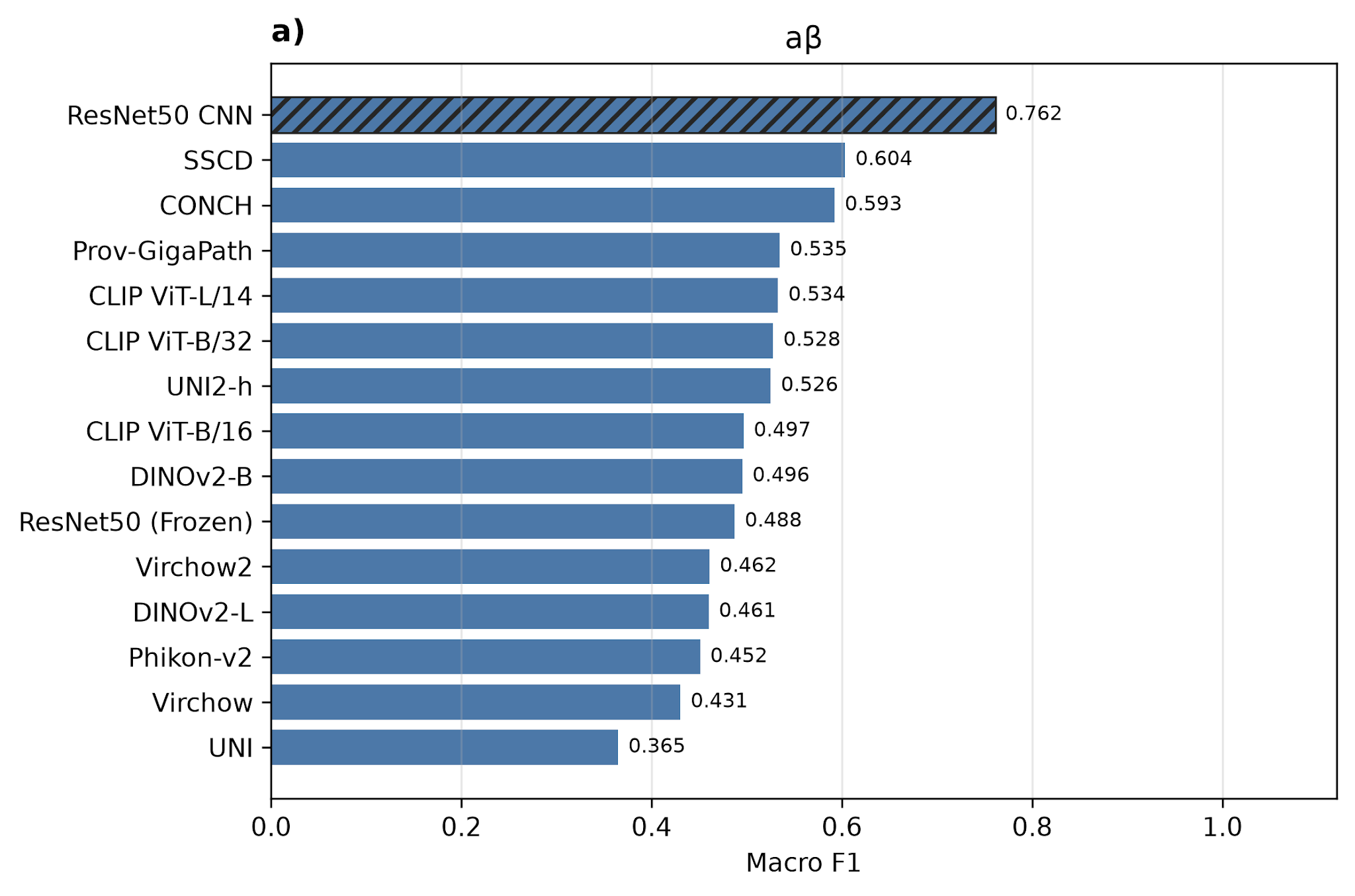

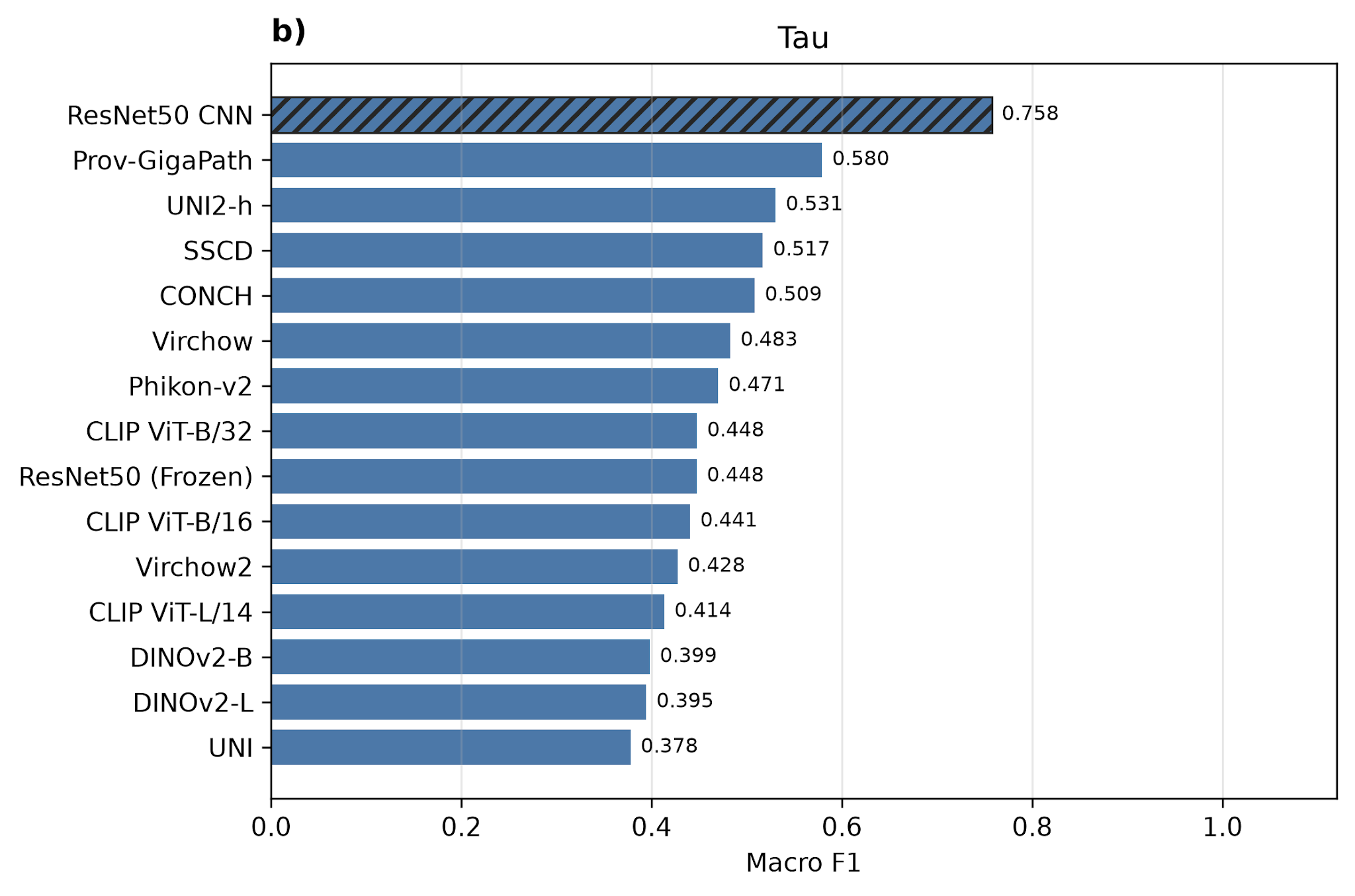

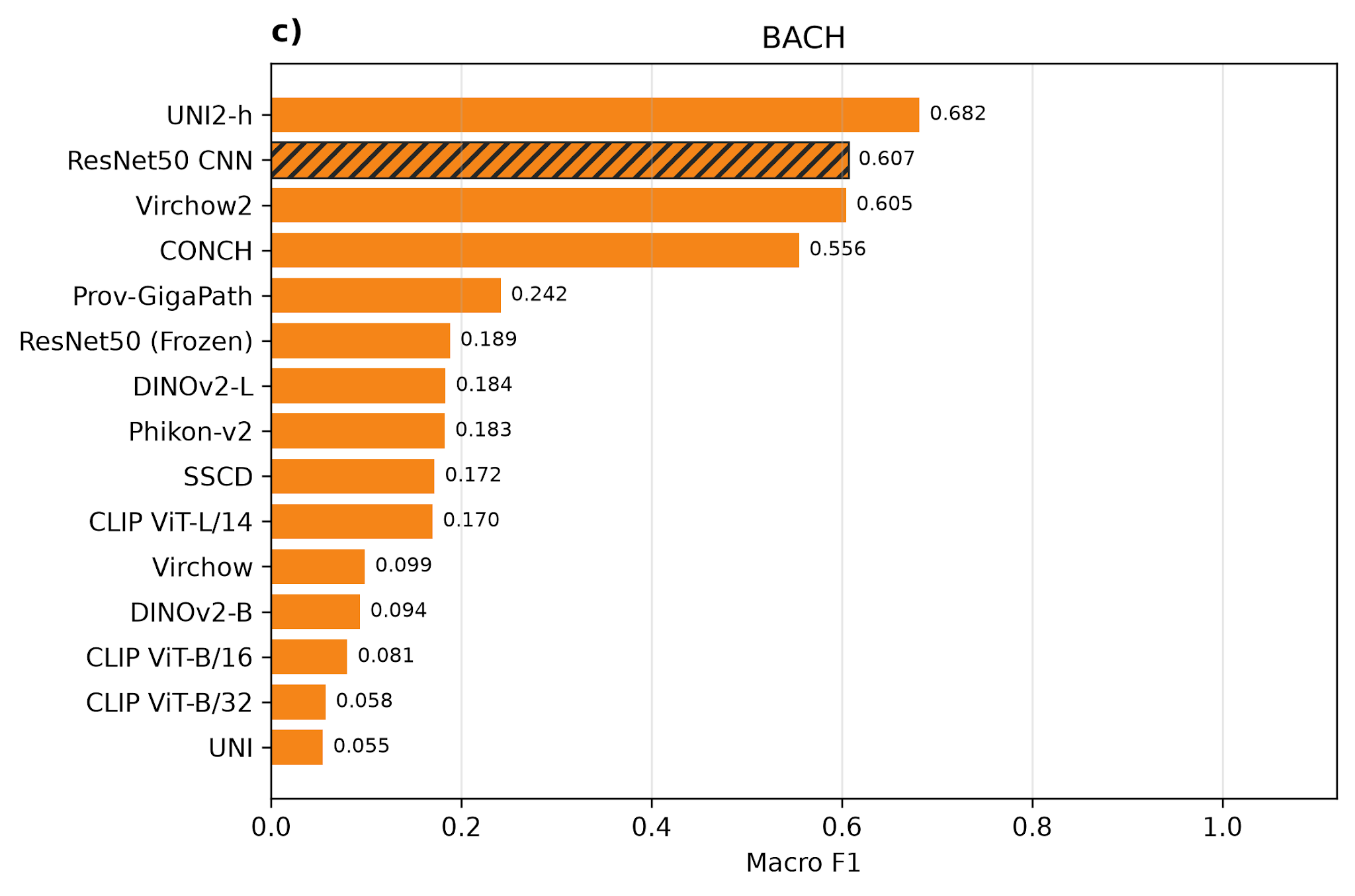

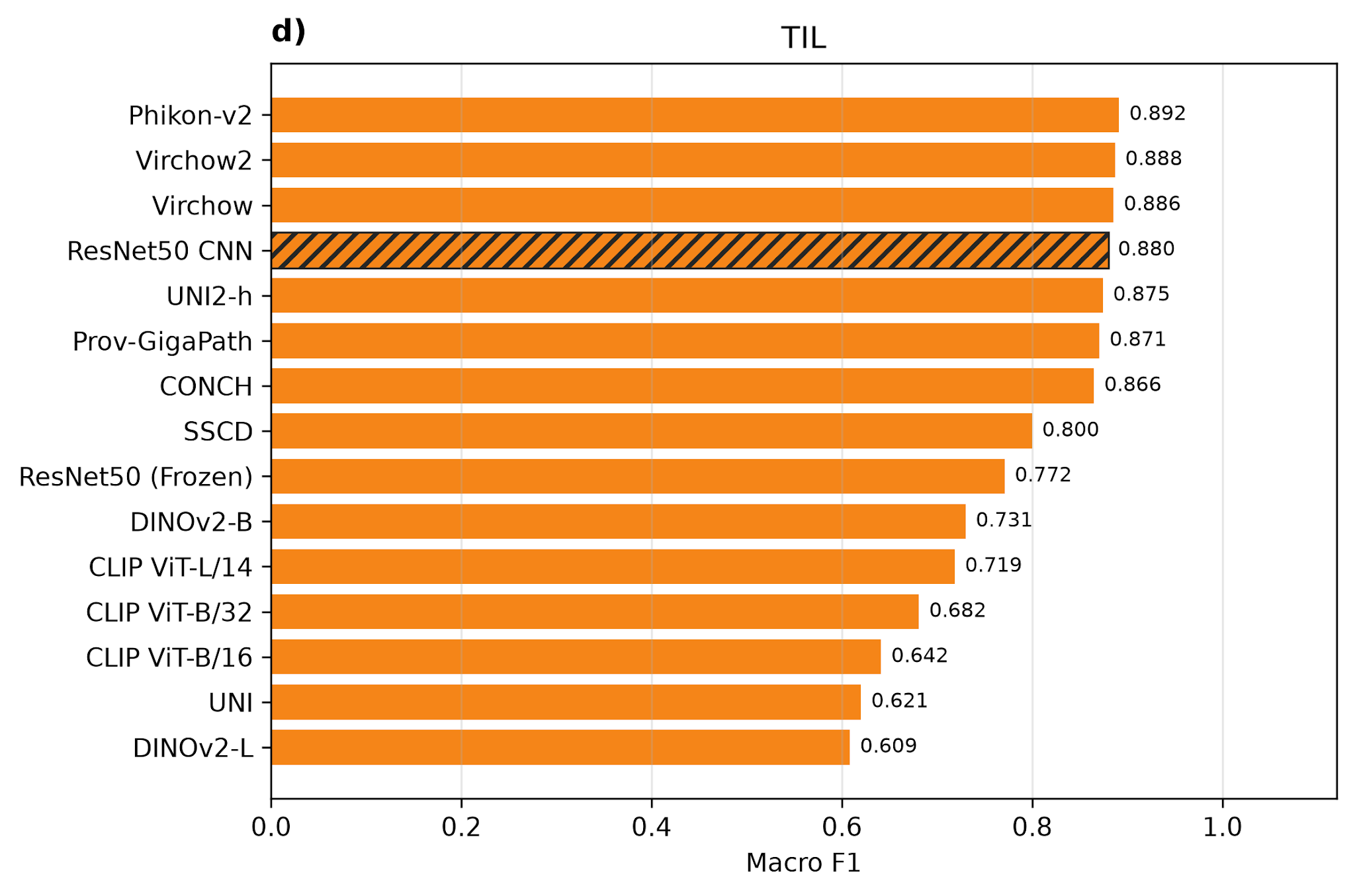

Supplementary Fig 1.Performance comparison of frozen foundation-model linear probes (solid bars) and a supervised ResNet-50 CNN (dashed bar) across four pathology datasets. Models are ranked within each dataset according to macro-F1 score. The CNN consistently achieved the highest performance on both neurodegenerative disease IHC datasets (a: aβ and b: Tau, blue bars), whereas pathology foundation models were competitive with or exceeded the CNN on the H&E datasets (c: BACH and d: TIL, orange bars).

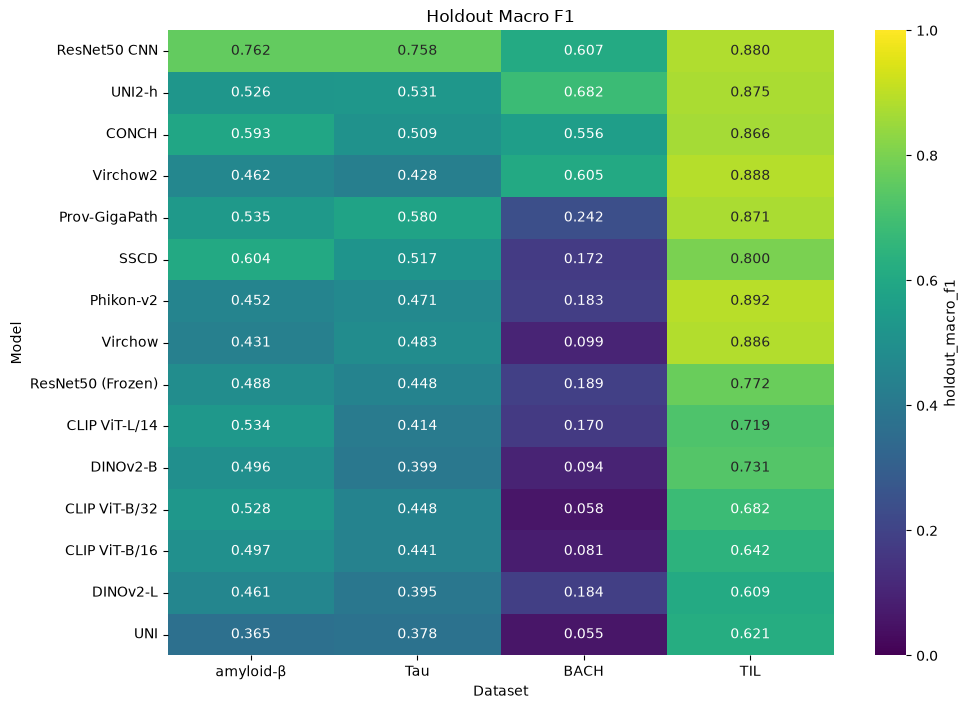
Supplementary Fig 2. Results of the various models on each of the datasets, shown in heatmap format for easy comparison.

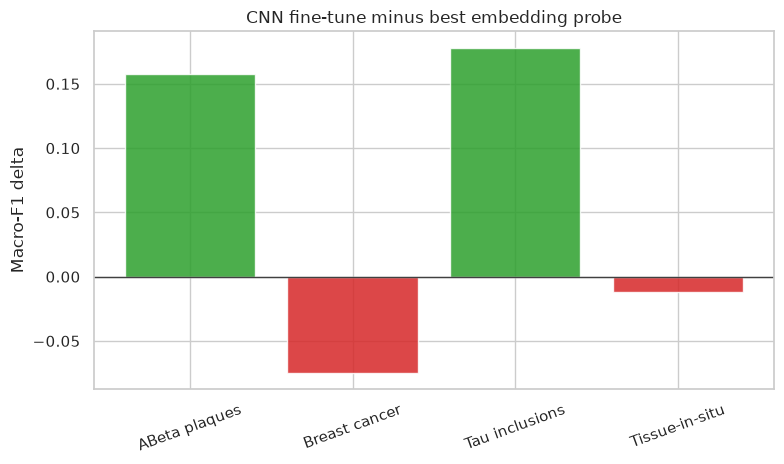

Supplementary Fig 3. The difference between the CNN and the best linear probe for each dataset, showing negative (red bars) when the linear probe is better and positive (green) when the CNN performs better.

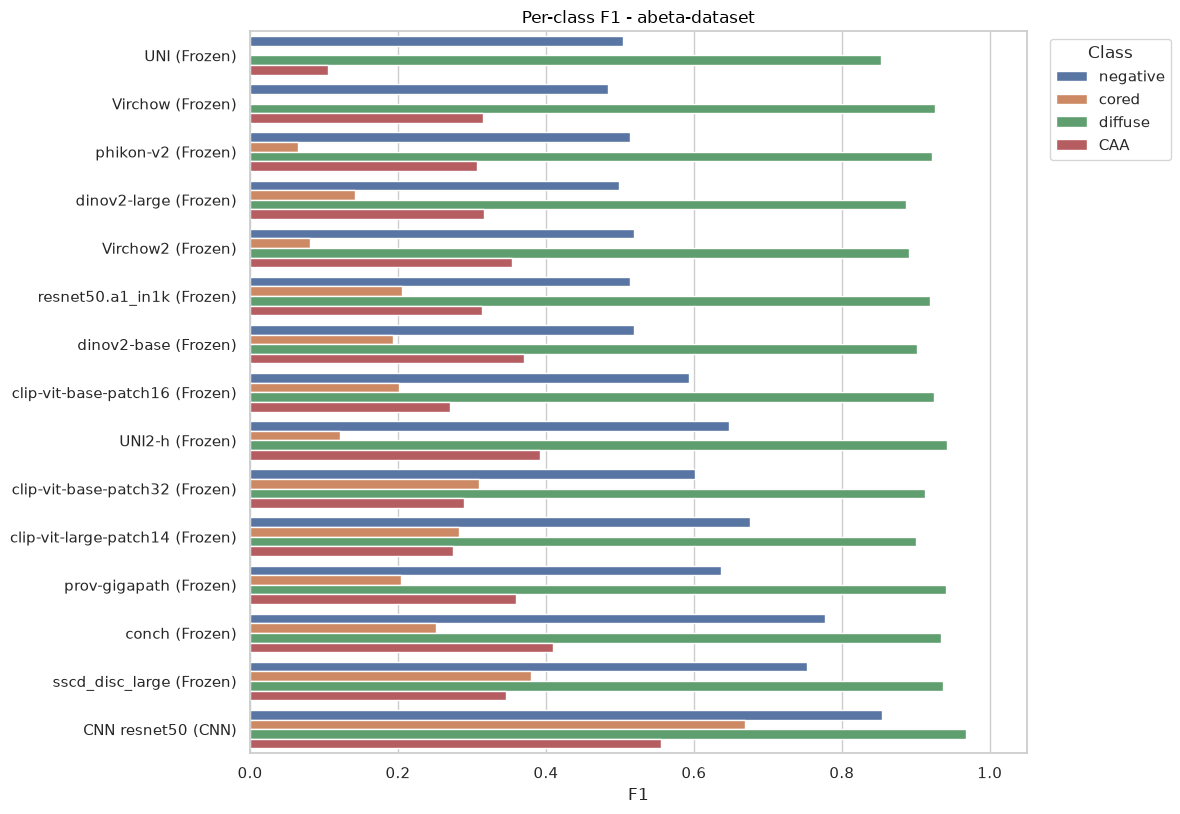

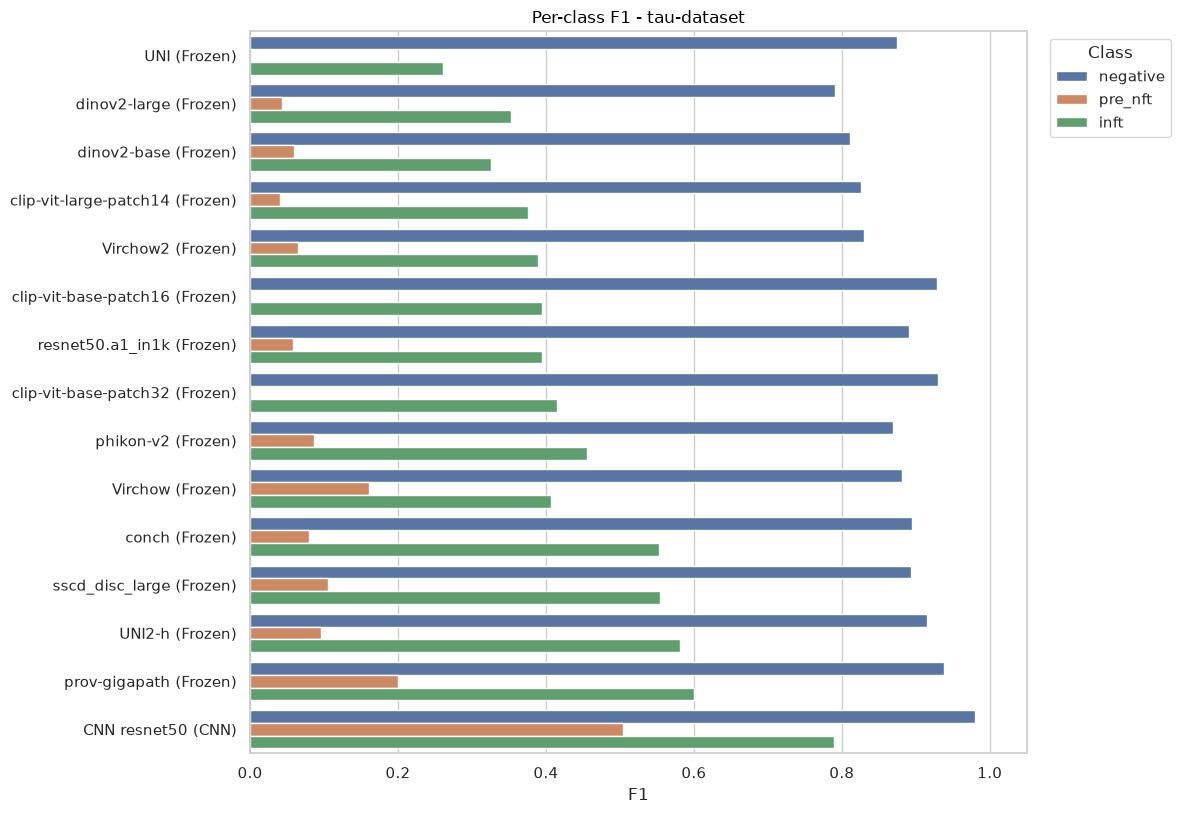

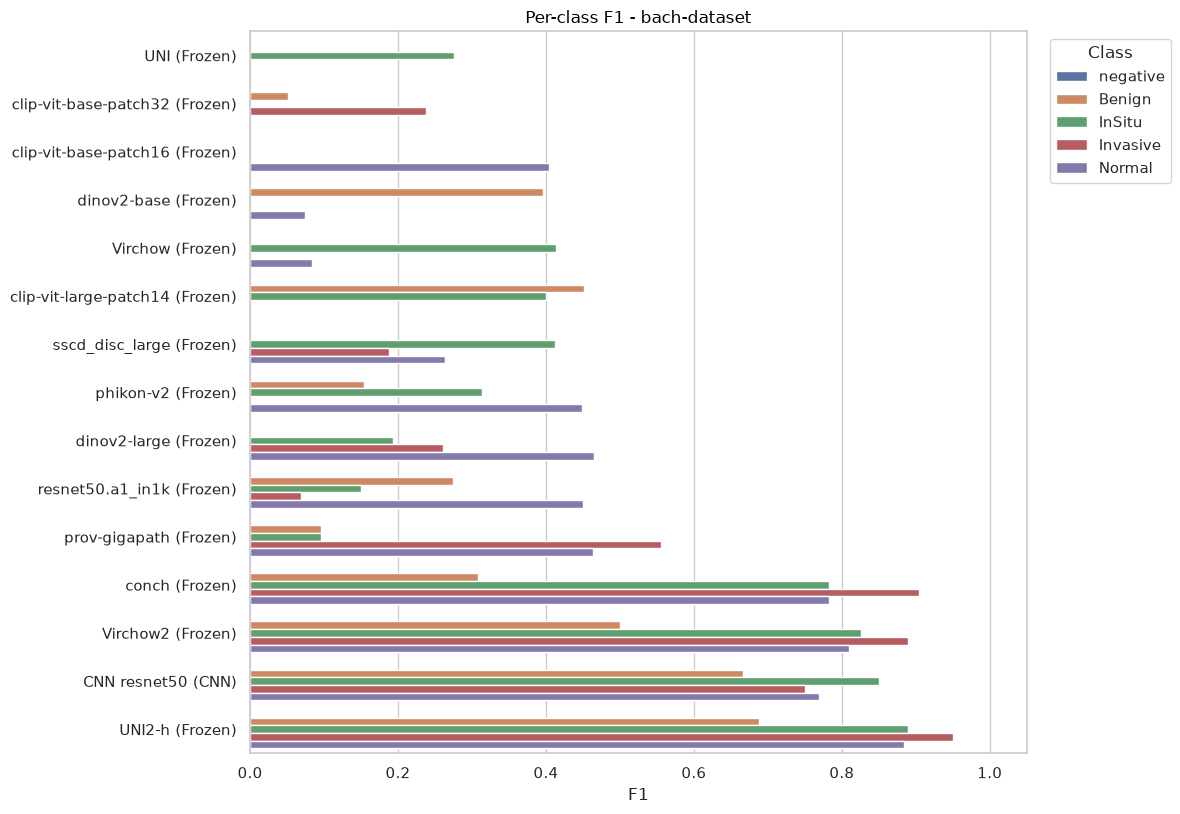

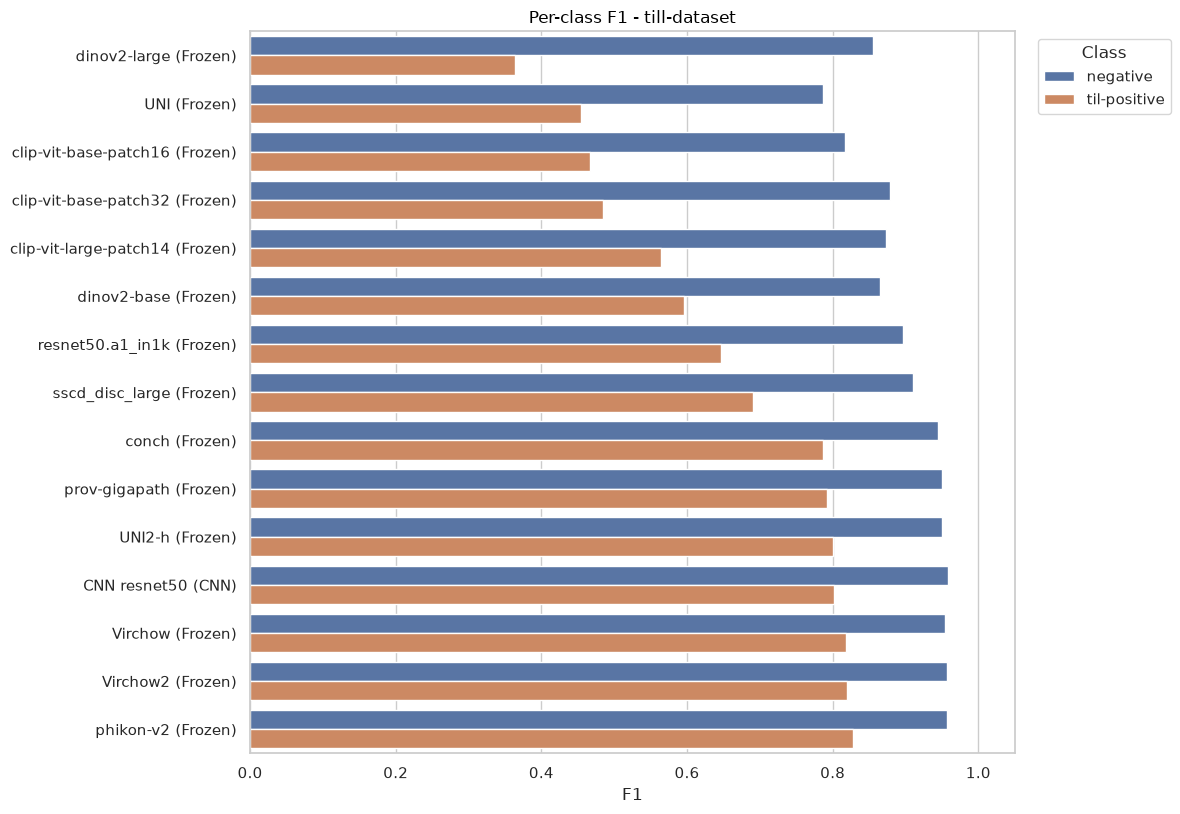

Supplementary Fig 4. The per-class F1 score for each of the datasets across all linear probes and the CNN.

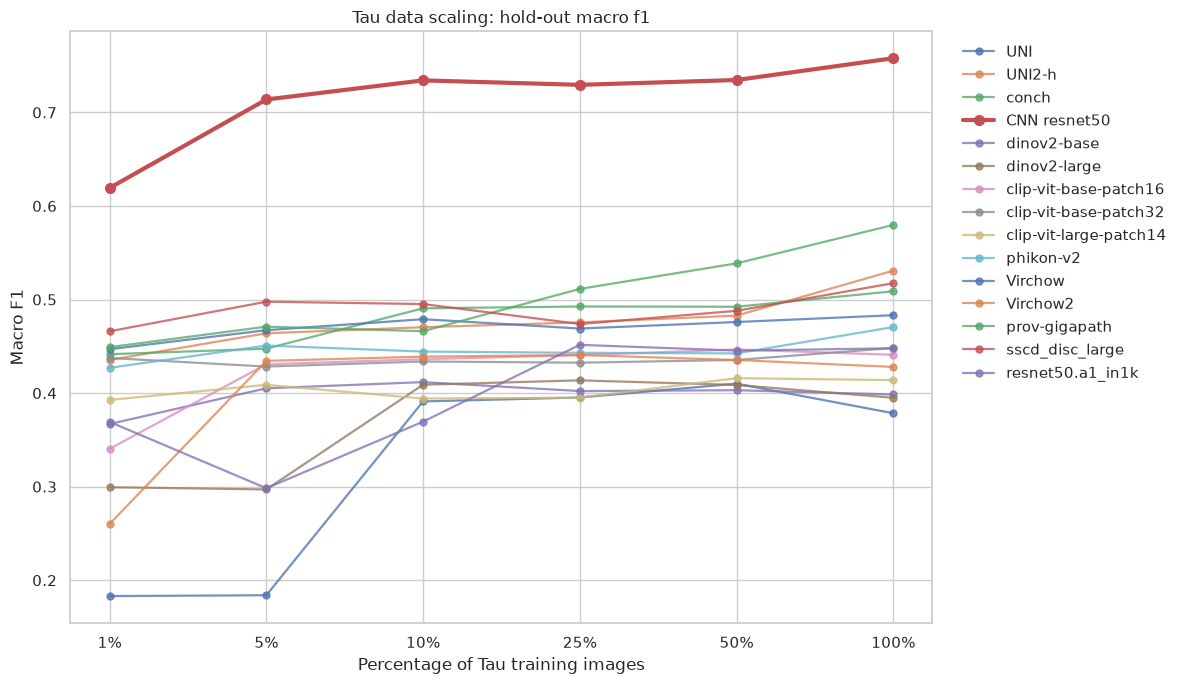

Supplementary Fig 5. The macro F1 score on the hold-out dataset for the scaling training data experiment. This figure includes the data shown in Figure 3 but also includes the rest of the models.
